# One More Lap: Environmental Modulation of Small-Scale Exploration in Weakly Electric Fish

**DOI:** 10.64898/2025.12.18.694964

**Authors:** Juan I. Vazquez, Valentina Gascue, Laura Quintana, Federico Pedraja, Adriana Migliaro

**Author notes:** Juan I. Vazquez. Now at: Department of Neurobiology, University of Konstanz, 78464 Konstanz; Germany; International Max Planck Research School for Quantitative Behaviour, Ecology and Evolution; Max Planck Institute of Animal Behavior, 78315 Radolfzell, Germany; Department of Collective Behavior, Max Planck Institute of Animal Behavior, 78315 Radolfzell, Germany. Valentina Gascue. Now at: Department of Biology, Boston University, USA.

## Abstract

Animals actively explore their surroundings to efficiently obtain resources, relying on the sensory modalities available to them. In South American weakly electric fish, self-generated electric organ discharges (EODs) create a short-range “sensory bubble” that, together with locomotion, supports exploration. This behavior relies on the coordinated modulation of spatial movement and electromotor activity, reflected in changes in EOD rate (EODr). However, how these components are engaged in natural contexts, and which conditions are sufficient to elicit exploration, remain poorly understood. Here, we examined how environmental and contextual factors modulate exploratory behavior in Gymnotus omarorum by combining a minimalistic laboratory assay with multi-day recordings in a seminatural arena. In laboratory tanks, freely moving fish showed minimal locomotor activity, which increased only when darkness and a novel electrosensory stimulus co-occurred, indicating strong context dependence and gating by chronobiological and motivational factors. By contrast, under seminatural conditions preserving natural light and temperature cycles, fish exhibited robust nocturnal rhythms in both locomotor activity and EODr. Chronobiological analysis revealed that the electrosensory rhythm consistently preceded the locomotor rhythm, with individual phase differences correlated across behaviors. These results show that exploration is a temporally organized behavior arising from coordinated modulation of sensory and motor systems. By linking electrosensory activity, locomotion, and circadian timing under ecologically relevant conditions, this study provides insight into how animals regulate sensory sampling and movement during exploration.

**Summary statement:** Exploration in *Gymnotus omarorum* is a temporally organized behavior arising from coordinated circadian modulation of electrosensory activity and locomotion.

## Introduction

Exploratory behavior is a fundamental trait for the efficient exploitation of an animal’s niche across taxa. It entails the acquisition of environmental information through an animal’s sensory toolbox, as well as physical movement through space to bypass the limits imposed by its sensory range. South American weakly electric fish gain information about their environment by constantly emitting discharges from an electric organ (electric organ discharge, EOD) which generates an electric field that surrounds the fish. A medullary pacemaker sets the frequency of EOD emission which is modulated by an array of environmental and physiological factors. Self-generated electric currents are in turn sensed by specialized sensory receptors embedded in the skin (Bullock et al., 1983; Caputi et al., 2005; Heiligenberg, 1977; Hopkins, 1974a; Hopkins, 1974b). Since the animal receives information only during the short duration of the EOD, exploratory behavior tends to be associated with increases in EOD rate (EODr). Locomotor activity also plays a crucial role. The body of the fish shapes the electric field due to a funneling effect that enhances perception in the mouth area where an electric fovea has been described (Caputi et al., 2022; Castelló et al., 2000). Despite being a remote active sense, the electrosensory system is a short-range system in the form of a sensory bubble (Caputi et al., 2013). Hence, exploration needs EOD emission as well as patterns of exploratory movements and locomotion to effectively align both the sensory epithelium and the sensory carrier towards objects in the environment (Hofmann et al., 2013). The fact that electric fish are nocturnal animals that inhabit highly turbid waters highlights the importance of the electrosensory system. In this sense there are several reports of nocturnal arousal as shown by increments in locomotor activity, EODr, EOD waveform and social interactions (Black-Cleworth, 1970; Silva et al., 2007; Stoddard et al., 2007; Zupanc et al., 2001).

South American weakly electric fish *Gymnotus omarorum* inhabit shallow turbid waters that are covered by thick vegetation patches, where adults establish 1m2 territories (Zubizarreta et al., 2020). Coincidently with their nocturnality, these electric fish show a nocturnal increase in EODr that follows a circadian rhythmicity. Hence the nocturnal increase persists in constant darkness in free running laboratory conditions, but also in the natural habitat, where sunlight is frequently blocked by the floating vegetation for long periods of time (Migliaro et al., 2018; Migliaro, Adriana, 2018). Locomotor activity has been less explored in this species. In a recent study, a remote sensing strategy allowed us to analyze electric and locomotor behavior in a natural undisturbed population that showed an increase in EODr and locomotor activity during the night as well as a strikingly high fidelity to the territory. Animals move more at night and resident animals patrol their territory and interact with an important number of visiting fish (Migliaro et al., 2025). Additionally, animals appear to prefer areas where light intensity is lower or darkness is permanent. In line with this, the exposure to light of natural intensities seems to be aversive, causing the animals to shelter and eliminating the characteristic EODr abrupt increases or bouts, thus reducing EODr variability (Camargo et al., 2023).

Exploratory behavior in electric fish has been approached in laboratory conditions considering both the electrosensory and the locomotor dimension. The novelty response, which entails a fast and short-lived increase in the EODr caused by the detection of sudden changes in the environment (Bullock, 1969; Heiligenberg and Bastian, 1980; Lissmann, 1958), has been thoroughly characterized (Barrio et al., 1991; Caputi et al., 2003; Grau and Bastian, 1986; Post and von der Emde, 1999). Together with this, fish might engage in “E-scans,” transient increments in EODr that anticipate the occurrence of a stimulus in locations where similar stimuli were sensed before (Jun et al., 2016). Locomotor patterns associated with electrolocation during exploration have been described in a variety of electric fish species as fish actively shape the acquisition of sensory information (Hofmann et al., 2013; Pedraja et al., 2020; Toerring and Moller, 1984). Movement reorients the sensory surface generating different complementary images of objects and exposing the sensory fovea to areas of interest (Castelló et al., 2000; Skeels et al., 2023). It also produces a sensory flow, increasing available dynamic information (Hofmann et al., 2013) and modifies the range and amplitude of the sensory carrier generated by the body of the fish, which is also a pre-receptor structure (Migliaro et al., 2005; Pedraja et al., 2014). In *G. omarorum* exploration has been assessed through novelty response in restrained animals, mainly by electrosensory (Caputi et al., 2003) and mechanical stimuli (Barrio et al., 1991). Reports associating EODr modulations with locomotor activity in the context of exploration are scarce. However, sustained (Falconi et al., 1995) and transient (Jun et al., 2014) modulations of EODr have been measured in association with locomotion and movement initiation.

Taken together, existing evidence indicates that exploratory behavior in weakly electric fish emerges from a dynamic integration of electrosensory sampling and locomotion. These behavioral components are embedded within a circadian framework and are therefore strongly shaped by environmental conditions. However, this integration has rarely been examined under experimental settings that allow for systematic manipulation of context while preserving ecological relevance in freely moving animals. *Gymnotus omarorum* operates within a complex sensory landscape structured by environmental cycles, spatial heterogeneity, and the presence of conspecifics and other organisms. Yet, the relative contribution of these factors, as well as the minimal conditions required to elicit and sustain exploratory patterns, remain poorly understood. In this study, we investigated how environmental and contextual factors modulate exploratory behavior, combining a minimalistic laboratory assay designed to identify stimuli sufficient to elicit exploration with multi-day recordings in a seminatural arena to assess the cyclic environmental modulation of this behavior.

## Methods

### Animal collection and housing

Specimens of *G. omarorum* (n=15) were collected at Laguna de los Cisnes, Maldonado-Uruguay (34°48’S, 55°18’W), using an electrical signals detector (Silva et al., 2003) and later housed in 500 l communal outdoor pools (1 x 1 x 0.5 m) in the animal facilities of IIBCE. Individuals were placed in perforated containers which prevented physical contact between neighbors while allowing exchange of electrical and chemical cues. Each container had a a PVC shelter. Pools were covered with plants obtained from the natural habitat, which included the free-floating water hyacinth *Eichhornia crassipes* and *Egeria densa*, *Myriophyllum aquaticum* and *Salvinia auriculata* (Zubizarreta et al., 2020). *Tubifex tubifex* was provided every five days. These conditions comply with national and institutional requirements (Ley N°18611, CEUA-IIBCE n° 001-02-2018). Animals were returned to communal pools after experiments.

### Laboratory experiments

Experimental set-up: fish (n=6) were individually placed in 48 x 38 cm (36.5 L) tanks inside a behavioral recording chamber (conductivity range: 60-200μS; room temperature: 25°C). This set-up has been used before to study both daily rhythmicity (Migliaro et al. 2016) and agonistic behavior in this species (Batista et al., 2012; Jalabert et al., 2015; Perrone and Silva, 2018). A 12:12 light/dark cycle was set to match the external facility environmental conditions (7:00 lights on, 19:00 lights off). Animals were allowed to acclimate to the experimental arena 24 hours prior to the experiment. During the experimental phase EODr and locomotor activity were continuously assessed from 17:00 to 11:00. An intermittent novel electrosensory stimulus was presented during 4 hours in two moments: from 17:00 to 21:00 (including when lights were off) and from 5:00 to 9:00 (including when lights were on). The stimulus consisted of a resistive object made from a plastic tube with conductive cylindrical carbon electrodes on both sides (dimensions: 5 x 10 mm, (Engelmann et al., 2009)) connected by a switch which was commanded by an Arduino board through Bonsai Rx (Lopes et al., 2015). Opening and closing the switch led to changes in the object’s resistance, which the animal was able to detect using their active electrosensory system. During the “stimulus on” period the interruptor was closed (20 Ω resistance) for 5s every 4min, completing 58 iterations. The sequential combination of both contextual factors determined six experimental conditions: stimulus-on/light at the evening (S/L_e_), stimulus-on/dark at the evening (S/D_e_), stimulus-off/dark (X/D), stimulus-on/dark at the morning before dawn (S/D_m_), stimulus-on/light at the morning (S/L_m_) and stimulus-off/light (X/L).

Electric recordings were made through a pair of copper electrodes attached to opposite sides of the tank. Signals were amplified (FLA-01, Cygnus tech) and digitalized through the computer’s motherboard soundcard commanded by Bonsai Rx, at a 10kHz sampling rate. Video was recorded by a chameleon3 (CM3-U3-13Y3M-CS) camera at 50 fps, with an infrared light, rendering 528 x 454 px videos. Video recordings were commanded by Bonsai rx and temporarily synchronized with electrical recordings.

EODr and locomotor activity assessment: EODs were recorded and the time of occurrence of every EOD was obtained using a threshold protocol on the z-score of the signal amplitude. EODr (expressed in Hz) was calculated as the inverse of the inter EOD interval. To study transient responses to the novel object a “response index” was created. This consisted in quantifying the appearance of high-frequency events (HFE, see supplementary material, figure S1), defined as instances in which the animals increased their EODr over 1.5 z-score for at least 3 consecutive EODs. The index was calculated as the percentage of instances of stimulus activation in which high-frequency events were observed. Baseline values for the presence of high-frequency events were calculated from random 10-second windows taken from recording periods when the object was not activated, for dark and light periods independently.

Locomotor behavior analysis was performed automatically using the open-source software DeepLabCut (DLC, (Mathis et al., 2018). A model was generated and trained (error after 4 training iterations, training: 0.28; testing: 0.40 cm). The model consisted of 5 points: mouth, head, pectoral fins, rostral end of the tail, and caudal end of the tail. Post-processing of DLC estimates was performed to smooth the tracking by erasing and linearly interpolating any prediction with a likelihood below 0.9. This allowed us to monitor locomotor activity in the whole area of the arena for the complete duration of the experiment.

### Seminatural Experiments

Experimental set-up: fish (n=9) were placed individually in 500 l outdoor pools, exposed to the natural environmental rhythms of light-dark and temperature cycles, as well as rainfall, with conductivity kept below 200 uS/cm and superficial vegetation brought from the natural habitat. To control that during the experiments seminatural conditions were within the expected range, light and temperature were recorded for each pool every 10 or 15 minutes (HOBO™ Pendant Temperature/Light Data Logger, https://www.onsetcomp.com/resources/documentation/ua-002-manual). Sensor ranges: temperature (-20 to 70 °C, thermic resolution and precision of 0.14 and 0.53 °C respectively); light (0 to 320,000 lux in a 100 to 1200 nm wavelength range). The daily thermal amplitude (calculated as the difference between mean daily maxima and minima) was 2,0°C ± 0,13°C. These measurements confirmed that our semi-natural setup reproduced light and thermal conditions typical of the species’ natural environment (Migliaro et al., 2005; Migliaro et al., 2018; Zubizarreta et al., 2020). EODs were recorded by eight electrodes (30 cm long, 5 mm diameter carbon rods) arranged in a 90 cm × 90 cm matrix. Electrodes were positioned from (x = 0, y = 90) in 45 cm steps, which allowed us to record across the whole area of the pool. Recordings were made by a signal acquisition system based on a multichannel amplifier controlled by a Teensy 4.1 board (Migliaro et al., 2005). This system allowed for continuous recording of the EODs. Recordings were stored as eight-channel WAV audio files of 1 min, with a sampling rate of 24kHz. Each animal was transferred to its experimental pool one day prior to the beginning of the experiment, for acclimation. EODs were recorded for six days EODr and locomotor activity assessment: Data was analyzed using *ad hoc* Phyton routines. For each individual the EOD data from the eight channels was squared and summed to enhance signal to noise ratio (see supplementary material for analysis routines). EODs were detected by a threshold protocol on the z-score filtered signal. Rate was calculated as the inverse of inter-event intervals. EODr was filtered to keep values between 6 and 70 Hz, which is well over the natural range of variation (Richer-de-Forges et al., 2009). Median rate and standard deviation were calculated for each 1 min file. EODr values were Q_10_ corrected to compensate for the direct temperature effects on metabolism (Ardanaz et al., 2001; Dunlap et al., 2000). Locomotor activity was calculated by changes in the amplitude of the EOD between electrodes. To obtain a movement parameter, the WAV files were segmented into consecutive 5-second windows. For each window, the median of the absolute amplitude of the signal was calculated for each channel. Based on these medians, the channel with the highest amplitude (dominant channel) was identified as an estimate of the fish’s position within the grid. A movement event was considered to have occurred when the dominant channel changed from one window to the next. To represent the probability of fish’ presence in the area, relative weights were assigned to three points: 1 for the electrode with the highest amplitude, 0.25 for the second highest, and 0.5 for the midpoint between them. These weights were accumulated for each spatial position across all windows of the file. Finally, the accumulated counts were normalized by dividing by the total recorded weight for each file, resulting in a proportional metric of space use.

### Data analysis

Statistical analysis was implemented in Python (libraries included in the supplementary material). Normality was assessed using the Shapiro-Wilk test. If normality was met, differences between experimental groups were evaluated using paired or independent t-tests. To assess the effect of experimental condition, moment of the day, and fish identity on locomotion RM-ANOVAs were used. In these cases, population data are presented as mean ± SEM. For data that were not normally distributed, Mann-Whitney U tests (for unpaired data), Wilcoxon tests (for paired data), and Friedman tests (with post-hoc Dunńs test) were applied. In these cases, data are presented as median ± confidence intervals (CI).

To assess the presence of daily rhythms in locomotor activity and EODr, a single-component cosinor analysis (Cornelissen, 2014) with a 24-hour period was applied using the Python library CosinorPy (Moškon, 2020). This library evaluates model significance through F and zero-amplitude tests (Bingham et al., 1982). Results are expressed as acrophases (in hours) and p values. All individuals (n=9) showed significant (p<0,05) daily rhythms for EODr while 8 individuals showed significant daily rhythms for locomotor activity. All analysis of locomotor activity was thus performed with n=8 individuals (Fig2) as well as comparisons between EODr and locomotor rhythmicity. Full cosinor parameters can be found in tables ST1 and ST2. To evaluate differences in individual acrophases, Rayleigh tests were applied to assess whether the phases were randomly distributed or showed significant clustering. Results are shown as Mean Resultant Length (MRL) and p values. To compare rhythmicity between locomotor and electric behavior we used linear regression when studying linear variables (R^2^), and Jammalamadaka-Sarma circular-circular correlations to compare acrophases. In this case to assess significance in mean acrophases we utilized the Watson-Williams test.

## Results

### Contextual modulation of exploratory behavior in a minimalistic arena

To gain knowledge on the contextual factors that promote exploration, we designed a minimalistic arena in laboratory tanks where animals were exposed to natural-like light/dark cycles and a single novel object to explore. Overall, animals showed extremely low levels of locomotion, spending under 3% of their time moving. In this setting, where the artificial light/dark context coincided with the natural day/night cycle, locomotor activity was minimal when the light was on if no novel object was presented to the fish. A global analysis that accounts for differences in locomotor activity of all the experimental conditions (light or dark, novel object present or absent) showed significant differences between contexts (Friedman’s Chi-squared test: F=15.686, p=0.0078). Post-hoc pair-wise comparisons showed differences in locomotor activity when both darkness and a novel stimulus were present compared to the lights-on/stimulus-off condition (Fig 2.A), both during the subjective evening and morning. However, the significance of these comparisons disappeared when correcting p values for multiple comparisons (Bonferroni-Holms correction, see supplementary tables 1 and 2 for the statistical summary). These results show that the co-occurrence of these two conditions is necessary to evoke locomotion. Regarding electric behavior, EODr showed no significant differences throughout the experiment and across experimental conditions. A reduced time window spanning from 6 to 8 pm (1 hour before to 1 hour after lights off) showed a transient EODr increment (see supplementary material), similar to what has been previously described for the same species in a similar set up without a novel object (Migliaro and Silva, 2016).

**Figure 1:**
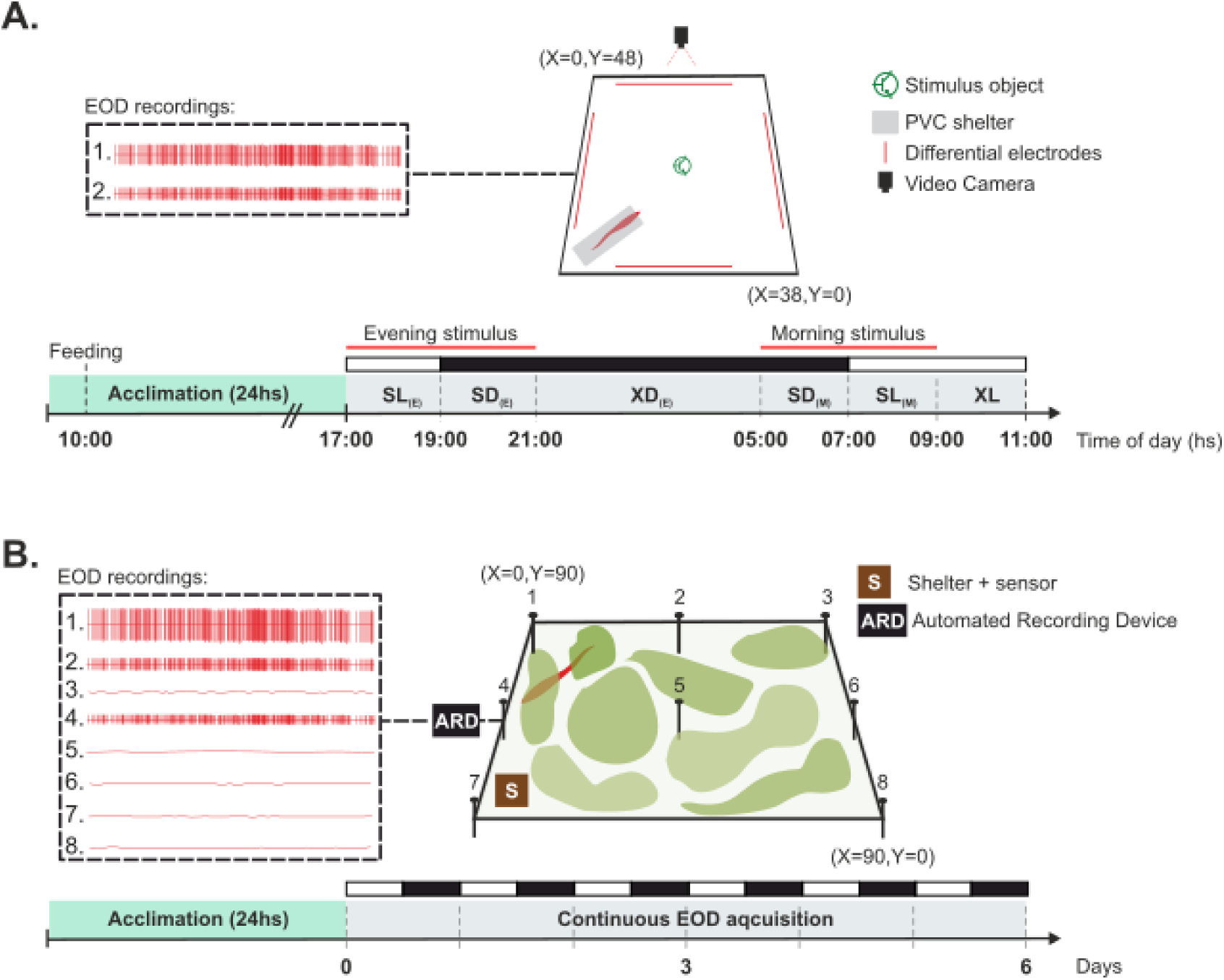
Experimental setup and recording protocols. **A.** Experimental design for the minimalist laboratory experiments. Left: representative EOD traces from the two pairs of differential electrodes. Middle: schematic of the experimental tank showing the PVC shelter, stimulus object, electrode placement (red), and video camera for behavioral monitoring. The bar below shows the experimental timeline. The white and black bars show when light was turned on and off respectively, and the red lines show the trial periods when the stimulus was turned on. In all, the experiment consisted of the following conditions: Stimulus on-Lights on during the evening (SL_E_), Stimulus on-Darkness during the evening (SD_E_), Stimulus off-Darkness (XD), Stimulus on-Darkness during the morning (SD_M_), Stimulus on-Lights on during the morning (SL_M_) and Stimulus off-Lights on (XL). **B.** Experimental design for long-term EOD acquisition in seminatural arenas. Left: representative EOD traces from different electrodes (1–8). Middle: schematic of the seminatural arena with differential electrodes (1–8) distributed across the arena, covered with natural vegetation. A PVC shelter equipped with a HOBO logger (S) was placed close to electrode 7. EODs were continuously recorded. The bar below shows the experimental timeline showing 24 h acclimation followed by 6 days of continuous EOD acquisition.

**Figure 2.**
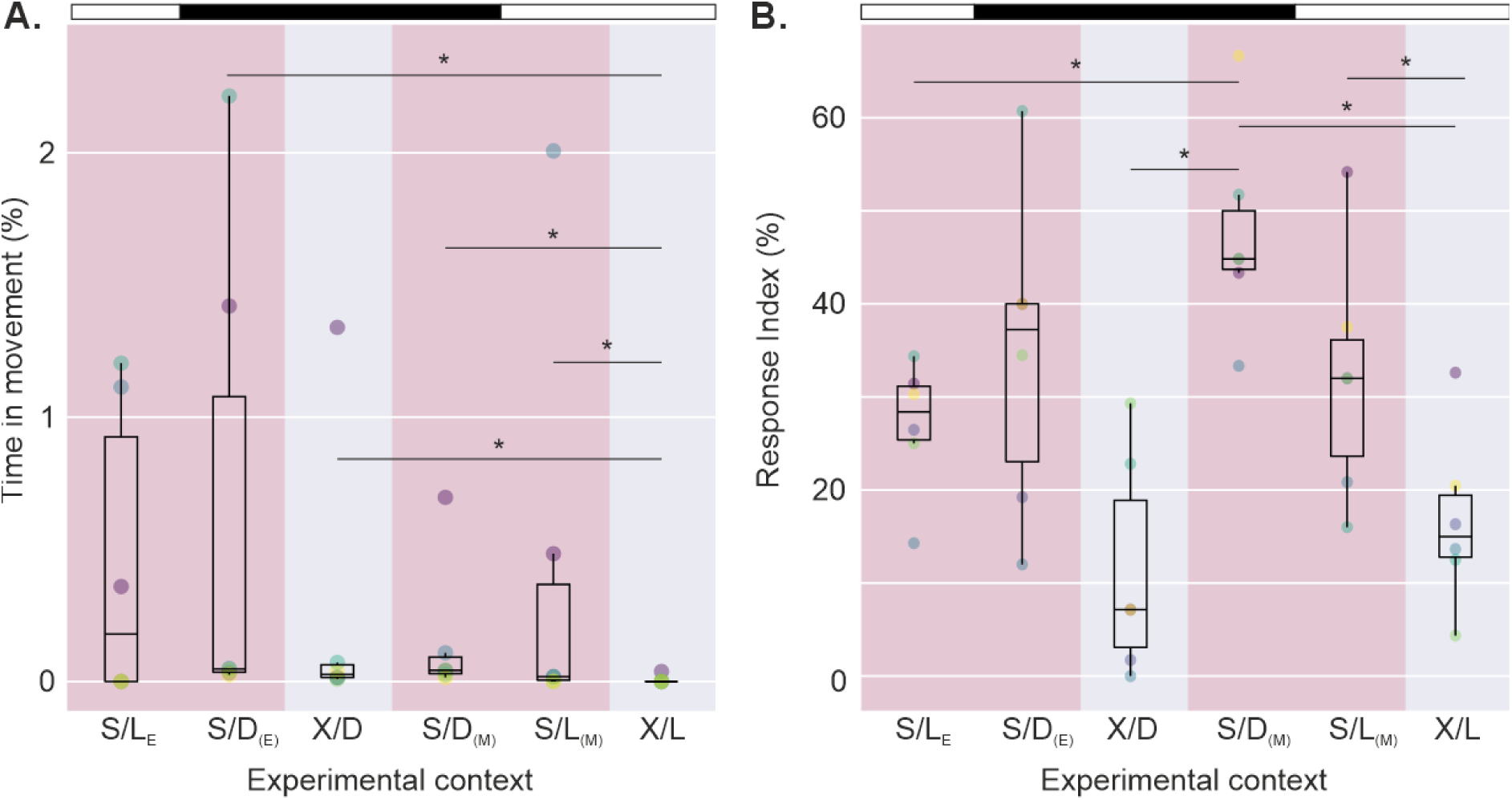
Changes in exploratory behavior under laboratory conditions when presented to a novel object. The white/black bar above the plots signals the light and dark phase respectively. The pink shaded area signals when the object stimulus was turned on. **A.** Percentage of time spent moving under different experimental contexts: Light or darkness (L/D) and with or without a novel stimulus (S/X) during the evening (E) or morning (M). Bars show median movement values with individual data points overlaid. Animals showed significantly different movement depending on illumination and the presence or absence of novelty (Friedman’s test: F=15.69, p<0.01). Asterisks mark post-hoc statistical tendencies (Wilcoxon’s signed rank test). **B.** Response index under the same conditions as A. Bars show median response index values with individual data points overlaid. Animals showed significantly different movement depending on the lighting and novelty contexts (Friedman’s test: F=19.81, p<0.01). Asterisks mark post-hoc statistical tendencies (Wilcoxon’s signed rank test), suggesting that both illumination and electrical stimuli influenced the amount of HFE.

Since resting EODr varied considerably among individuals (from 33,3 to 42,7 Hz) and considering that animals that had higher resting EODr might need a smaller increase in their rate to electrically explore an object, we tested for a possible correlation between the response index and the resting EODr using linear regression and did not find a significant relationship (R^2^= 0.015, p=0.815, not shown). The electrosensory response index showed significant differences between contexts (Figure 2B, Friedman’s Chi-squared test: F=19.81, p<0.01). Multiple comparisons analysis shows that this index was significantly higher when lights were off and the object was being intermittently turned on. As with locomotor activity, this significance disappeared when correcting for multiple-comparisons (see table S2 for full summary). The small number of animals (n=6), the relatively low amount of movement (less than 3% of the time) and the interindividual variability in the resting EODr might explain the reduced statistical power to detect pairwise differences. However, our data confirms a global effect of illumination and novel stimuli on the modulation of exploratory behavior.

### Contextual modulation of exploratory behavior in a seminatural arena

Locomotor activity was measured throughout six consecutive days in isolated individuals placed in experimental pools. Fish show a clear daily pattern with higher amounts of movement during the night. Well differentiated resting versus active states become evident from this analysis. Figure 3A shows the median locomotor activity throughout the day for each fish (colored lines), as well as the population-wide median value at every hour (black line). A cosinor based analysis confirms a 24-hour rhythm for all animals (see complete cosinor parameters in the supplementary material). A global analysis of this daily locomotor rhythm (fig 3B) shows that maximum locomotor activity is reached towards midnight, with a mean population acrophase at 00:33 hs (Rayleigh test: MRL=0.86; p<0.001). The average MESOR was 11.9% ± 1.78%, while the amplitude of the rhythm was 7.95% ± 1.46%.

**Figure 3.**
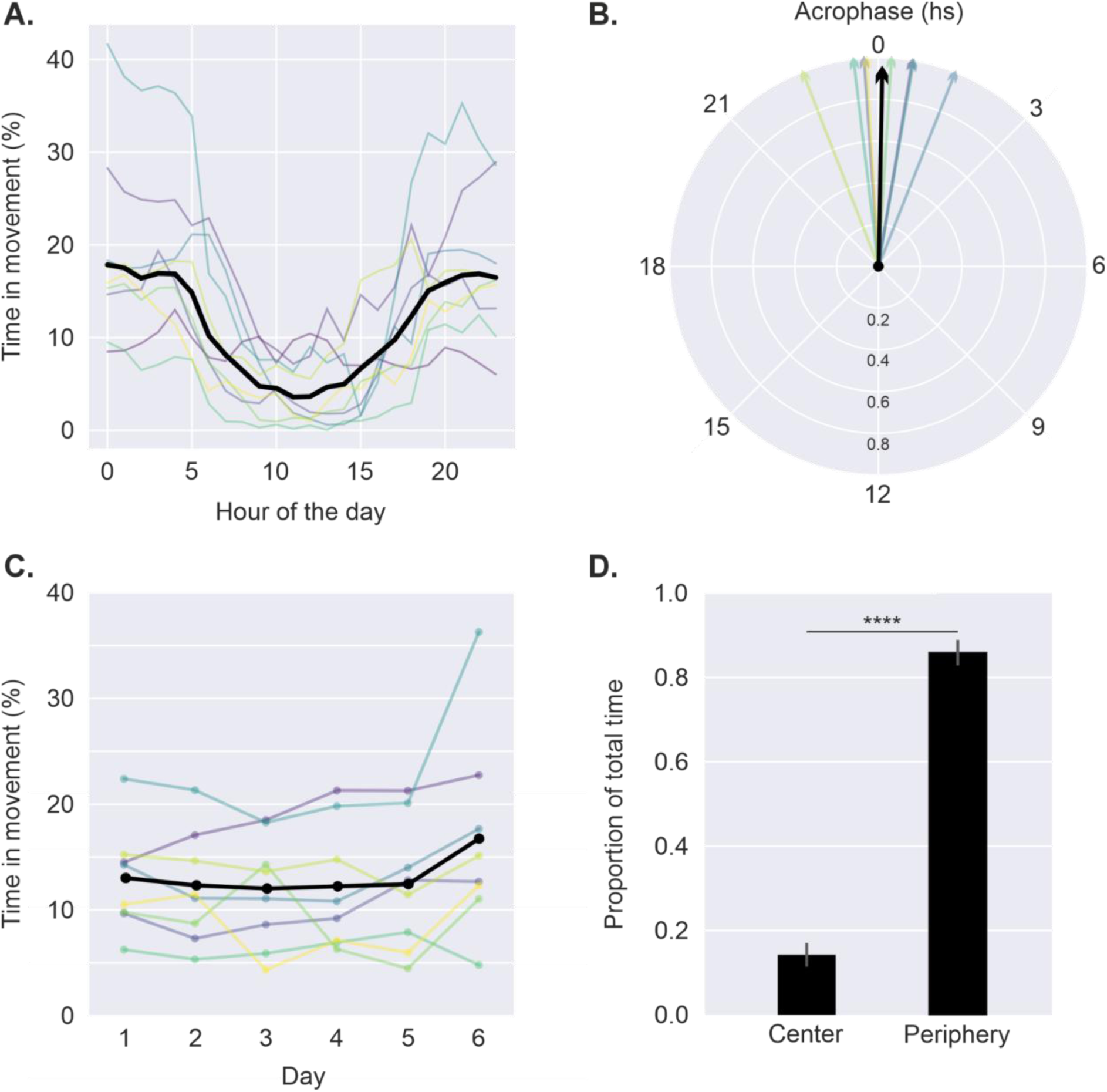
Daily locomotor activity and spatial distribution of fish under seminatural conditions. **A.** Mean locomotor activity across the day. Thin colored lines represent individual fish while the thick black line shows the population mean. **B.** Polar plot showing the acrophases obtained through cosinor analysis for individual fish (color arrows) as well as the mean acrophase (black arrow). The analysis shows clear directionality between acrophases (Rayleigh test: MRL=0.86; p<0.001). **C.** Locomotor activity across experimental days. Thin colored lines indicate individuals, and the black line shows the population mean (see text for statistical data). **D.** Proportion of total time spent in the center vs. periphery of the tank. Bars represent mean ± SEM. Animals spent significantly more time using the periphery of the arena (paired t-test: p < 0.0001).

In order to rule out a decline in locomotor activity due to captivity, the percentage of time in movement was calculated for each of the six days (Fig 3C). An RM-ANOVA analysis confirmed differences in average movement between days (RM-ANOVA: F (5, 35) = 2.81; p <0.05). However, a post-hoc analysis revealed that this difference was just due to an increase in activity values on the 6th day compared to the first day that only reached trending significance (paired t-test: p=0.07).

The precise location of the electrode array allowed us to quantify how much time animals spent using different regions of the seminatural territory. Regardless of the moment of the day animals tended to remain in the periphery of the arena even though the water surface was partially covered with natural vegetation that might have offered alternative shelters and multiple food locations (Figure 3D).

As previously reported for this species in the natural habitat, electric behavior shows a clear daily pattern of EODr. Figure 4A shows individual median hourly EODr and the population median values. Animals reach minimal EODr values early in the morning gradually increasing towards the evening. As with locomotor activity a cosinor based analysis confirms a 24-hour rhythm for all animals in this experiment (see complete cosinor parameters in the supplementary material). Individual acrophases tend to accumulate early in the night (Fig 4B). A global analysis shows a mean population acrophase at 21:49 (Rayleigh test: MRL=0.86; p<0.001). As a further validation of our method, we studied if the EODr levels decreased as the experimental days advanced (Fig.4C), which would indicate welfare problems in the individuals. In this regard, we found no relationship between the median EODr and the experimental day (RM-ANOVA: F (5,35) = 1.78, p=0.1423).

**Figure 4:**
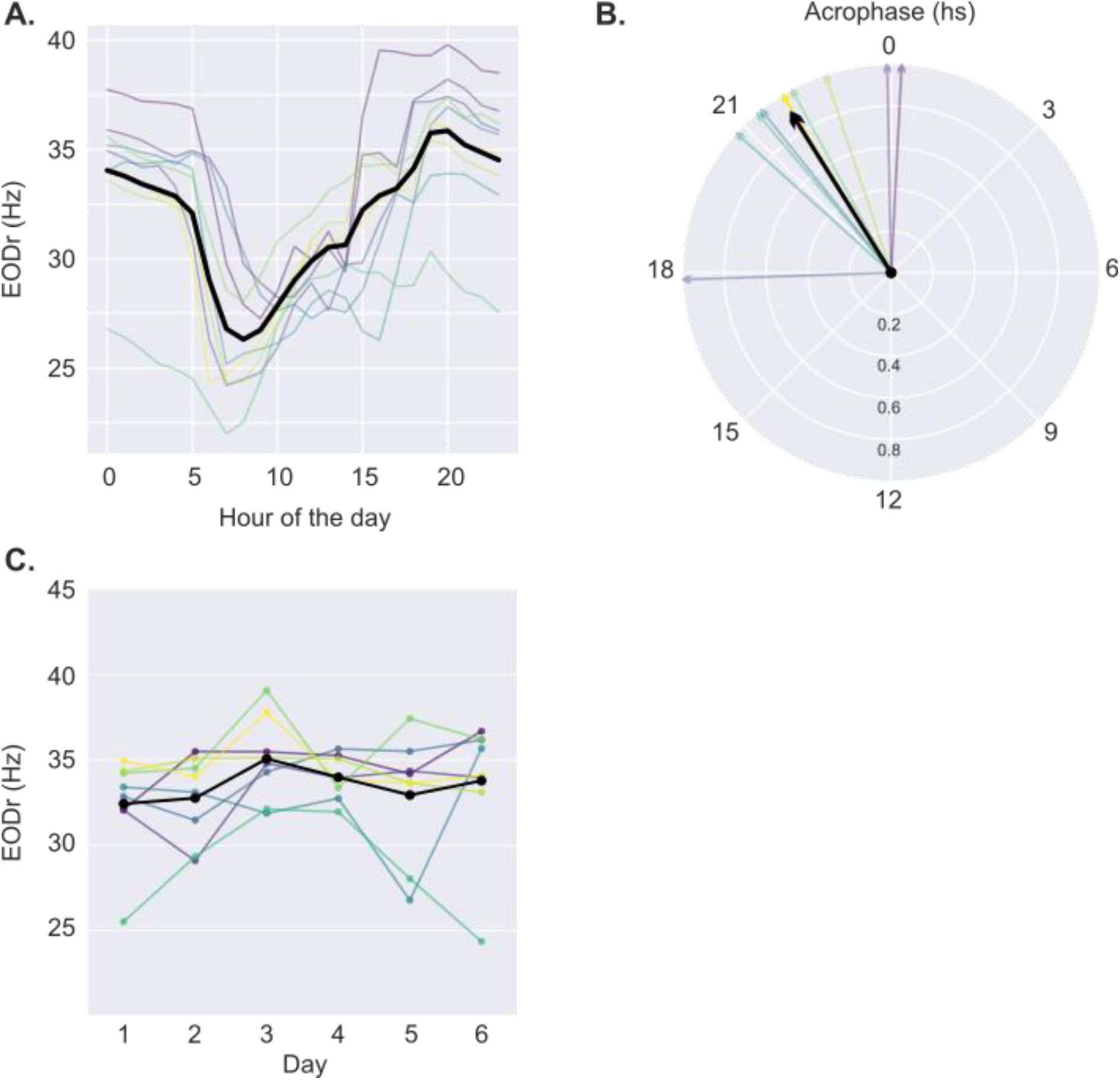
Daily rhythm of EOD rate in *Gymnotus omarorum* under seminatural conditions. **A.** Mean hourly values of EOD rate across individual fish (thin colored lines) and median population values (black line). EODr shows a clear circadian modulation, with a trough during the morning and a peak in the evening. **B.** Polar plot showing the acrophase of EODr rhythms estimated through cosinor analysis for each fish (colored lines). The black arrow shows the population mean acrophase. The length of the arrow corresponds with the MRL of the individual acrophases. Acrophases show a clear directionality (Rayleigh test: MRL=0.86; p<0.001). **C.** Median EODr values across experimental days. Thin colored lines indicate individuals, and the black line shows the population mean. EODr values remained stable throughout all experimental days (RM-ANOVA: F (5,35) = 1.78, p=0.1423).

We analyzed the temporal relationship between the locomotor and electrosensory components of exploratory behavior. Electrosensory rhythm acrophase anticipates locomotor acrophase by approximately 3 hours (Fig 5A, Watson-Williams test, p<0.01). In order to assess whether the electric and locomotor behavior followed a concordant temporal organization, circular-circular correlations between EODr and locomotor activity acrophases were obtained for each fish. A strong correlation between individual acrophases of both behaviors shows that animals with an earlier pattern in electrosensory rhythm will express an earlier locomotor rhythm (Fig 5B, Jammalamadaka-Sarma circular-circular correlation: ρ=0.75, p<0.05). No correlation was found between the robustness of each rhythm measured by the R^2^ parameter (Fig. 5C, p=0.97), and electrosensory rhythms are more robust than locomotor rhythms (Fig 5D, paired t-test: p<0.05).

**Figure 5:**
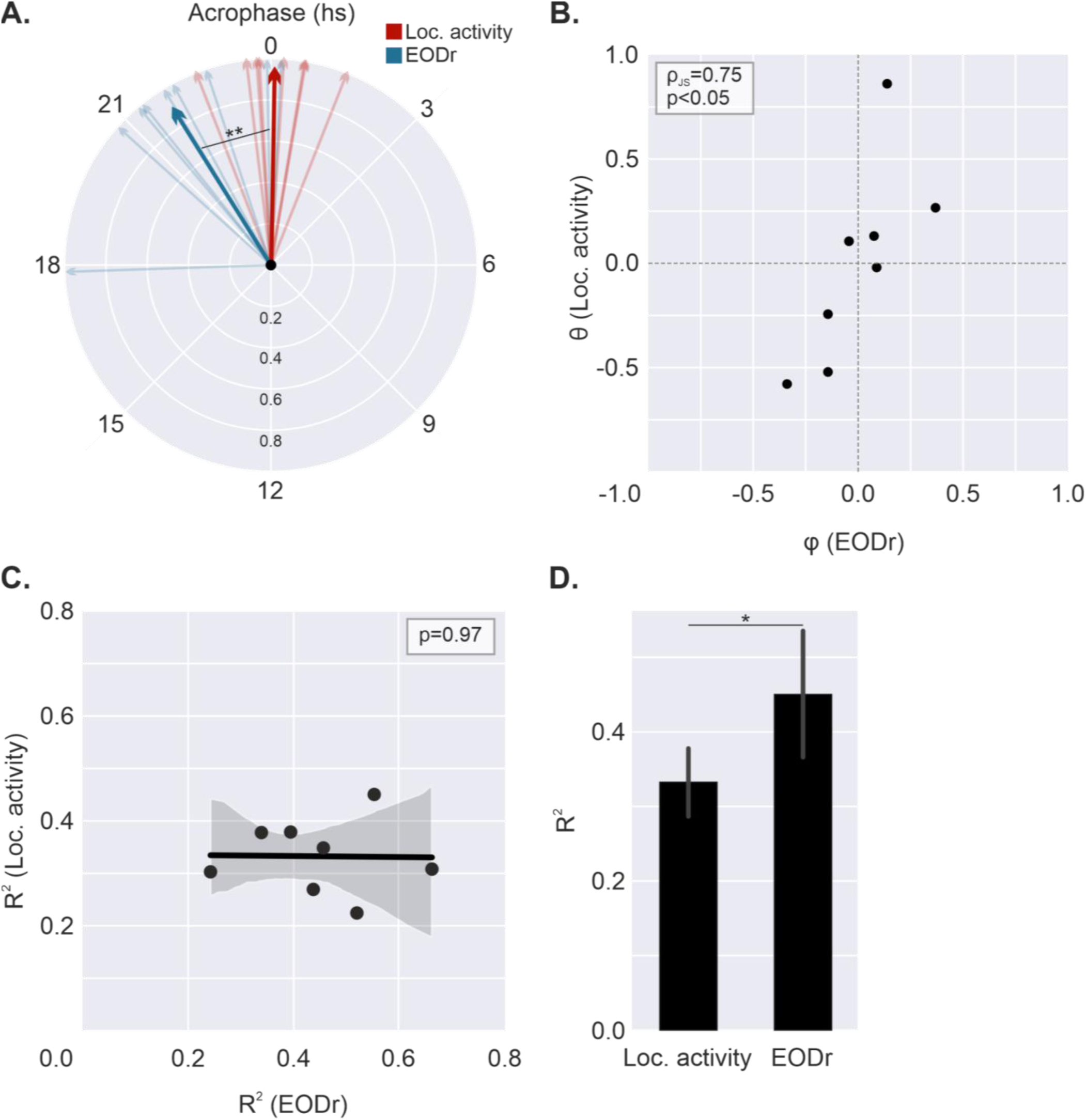
Comparison of rhythmic parameters between locomotor activity and EODr. **A.** Polar plot showing the individual and population acrophases for both the locomotor (red) and EODr (blue) rhythms. The mean locomotor acrophase was significantly later than that of the EODr rhythm (Watson-Williams test: p<0.01). **B.** Circular correlation between acrophases of locomotor activity (θ) and EODr (φ). A significant positive correlation was detected (ρ_JS_ = 0.75, p < 0.05), indicating concordant alignment of rhythms across individuals. **C.** Linear regression between the percentage of variance explained by the cosinor model (R²) in locomotor activity and EODr across individuals. No significant relationship was found (p= 0.97). **D.** Mean R² values (± SEM) for locomotor activity and EODr. Rhythmicity was significantly stronger in EODr compared to locomotor activity (paired t-test: p < 0.05).

## Discussion

*Gymnotus omarorum* displays a robust nocturnal pattern of exploratory behavior, integrating both locomotor and electrosensory components. Exploratory behavior is fundamental for exploiting the resources of a territory. Weakly electric fish explore their surroundings combining electrolocation and movement. EODs provide the basis for electric image formation and several parameters of this image encode the various features of objects in a scene (Caputi and Budelli, 2006). Since EODs constitute the carrier of electrosensory stimuli any increase in the rate of emission renders an increase in the amount of sensory information available. Locomotor behavior is also crucial for exploratory behavior. On the one hand considering that electrolocation is a short range sensory system spanning a couple of centimeters around the fish body, displacement is crucial for spatial exploration, “carrying” the self generated electric field around the territory. On the other hand, the fish body acting as a pre-receptor structure modifies the electric field by adopting different postures which also differently expose the sensory epithelium (i.e. the skin) enhancing perception. In previous work (Migliaro et al., 2025) we studied locomotor patterns of an undisturbed population of this species. This approach was able to capture that EODr and locomotor activity were higher during the night, that animals exhibit remarkable site fidelity, and that the occurrence of apparent social interactions between two fish resulted in one of them leaving said territory. However, the spatial resolution and time-scale did not allow us to explore how animals were actively using their territory or to keep track of individual behavior for long periods of time.

In this study, we examined the expression of exploratory activity in *G. omarorum* through its locomotor and electric behavior under two complementary conditions. The first consisted of a minimalistic, plain-water arena with a controlled 12:12 light–dark cycle and a single novel object, designed to identify the minimal conditions sufficient to elicit exploration. We found that both darkness and novelty promoted exploratory behavior, though they acted differently on locomotor and electric components. The second approach consisted of a seminatural arena that mimicked a simplified territory—free of conspecifics and predators but maintaining territory size, natural light and temperature cycles, surface vegetation, and associated invertebrates. Exploration in this context likely reflects the combined influence of environmental rhythms, foraging, and territorial patrolling. Under these seminatural conditions, fish exhibited stable and well-defined nocturnal rhythms in locomotion and EODr, with the moment of maximal electrosensory activity preceding the peaks of locomotor activity. This temporal coordination likely optimizes environmental sampling under natural cycles. While during the night, *G. omarorum* will actively explore a seminatural enclosure that mimics the natural territory, in the minimal laboratory setting exploration was strongly context-dependent and co-modulated by darkness and novelty.

### Exploratory behavior in the minimalistic arena

Exploratory behavior is shaped by environmental and contextual factors, particularly light and novelty. In this study, a significant overall effect of condition was observed, suggesting that exploratory behavior changes systematically with these contextual modulators. Darkness and the presence of a novel object acted synergistically to enhance movement, emphasizing the inhibitory role of light—consistent with the species’ nocturnal ecology and previous observations in both field and laboratory settings (Camargo et al., 2023; Migliaro et al., 2025). Although overall locomotor activity was markedly reduced, it increased significantly when both permissive (dark) and stimulating (novel) conditions co-occurred. Electrosensory activity, by contrast, showed a transient rise in EOD frequency at lights-off, similar to earlier reports (Migliaro and Silva, 2016), and was more strongly influenced by novelty than by illumination. These results underscore that information on the immediate surroundings modulate exploratory behavior at multiple levels: novelty triggers exploration, while nocturnality enhances its expression. Together with these light acts as a suppressive signal for locomotor activity. The absence of exploratory activity in the laboratory, unless both chronobiological (darkness) and motivating (novelty) conditions were present, further supports the hypothesis that *G. omarorum* regulates exploration through the interaction between environmental context and endogenous rhythms as well as the internal motivational state. Although darkness and novelty significantly modulated exploratory behavior, post hoc comparisons did not remain significant after Holm–Bonferroni correction neither for locomotor activity nor for electric behavior. This loss of significance is likely related to the combination of a small sample size (n = 6), the small percentage of time in movement, and interindividual variability in EOD basal rate. These factors reduce the statistical power to detect pairwise differences, even in the presence of a consistent overall trend. Therefore, the results should be interpreted with caution: although the data point toward a potentially relevant contextual effect, additional studies with larger samples are needed to confirm and better characterize these differences.

### Exploratory behavior in the seminatural arena

The close correspondence between the daily rhythms of EODr and locomotor activity supports the idea that exploratory behavior in weakly electric fish involves a dynamic interplay between sensory acquisition and movement. Previous work has shown that EODr exhibits circadian modulation in *G. omarorum* (Migliaro et al., 2018) and other gymnotiform species (Dunlap et al., 2011; Stoddard et al., 2007). Our results extend these observations by demonstrating that locomotor activity follows a comparable nocturnal pattern. Our data shows that both EODr and locomotor activity increase at night, reflecting the nocturnal nature of the species. Both traits were known to increase in nature for freely moving animals, however, this experimental design has allowed us to keep track of known isolated individuals for several days, which is mandatory for the study of circadian rhythms. Daily rhythmicity was assessed by the cosinor model for each animal. Rayleigh tests for each behavior confirms a coherent rhythm of the whole population. These animals, kept in isolation but subjected to similar environmental fluctuations, express a synchronized rhythmicity that reflects the influence of a cyclic environment on the individual endogenous rhythms. A comparison of the locomotor and electric rhythmicity shows that electric behavior reaches maximum values nearly 3 hours earlier than locomotor activity. In fact, a strong positive correlation exists between the phases of these rhythms, underscoring the fact that the nocturnal increase in electric behavior always precedes the nocturnal increase in locomotor activity, and that an early EODr rhythm is associated with an early locomotor rhythm. Individual analysis of biological rhythms in humans has led to the concept of chronotype: the identification of individual differences in the timing of daily activities (Horne and Ostberg, 1976). The abstract idea of an individual preference of a certain phase difference between behavioral rhythms and the day/night cycle is strongly correlated to physiological indicators of circadian phase (Roenneberg et al., 2003; Tassino and Silva, 2024). In this sense and for the same environmental cycle, rhythms have earlier acrophases in individuals with early chronotypes and vice versa. More recently this concept has been extended and adapted to non-human animals considering chronotype as timing of activity onset relative to environmental cues (Maury et al., 2020). This opens a promising path to link individual variations in the phase-locking to environmental cues to a wide variety of genetic, neural, physiological and behavioral scenarios (Graham et al., 2017; Maury et al., 2020; Meijdam et al., 2022; Tomotani et al., 2024). In this sense our data allow us to identify early explorers and late explorers much in the well-known lark/owl classification, using either EODr or locomotor acrophases as a proxy for chronotype. Future expansion of research on wild chronotypes is extremely promising in the quest for understanding how the brain generates efficient behavior which is timely adapted to the natural environment, particularly considering natural individual variability.

The relationship between these two components of exploratory behavior can be analyzed at different temporal scales. Laboratory recordings have shown that locomotor events are often preceded by transient increases in EODr that have been classified as evidence of volition (Jun et al., 2014). Likewise, increases in EODr are associated with spatial recognition and with food searching or predictive exploration in places where electrosensory stimuli had previously been detected (Jun et al., 2016). On the other hand, the daily rhythm of EODr is largely independent of locomotion, since animals which are voluntarily motionless also display this characteristic nocturnal increase (Migliaro et al., 2018). Within the daily timescale relevant to this work, we observed that the rhythm of electric behavior precedes the locomotor rhythm. Such coordination may optimize the use of energy by enhancing sensory vigilance during periods when locomotion — and thus predation risk or energy expenditure — is still minimal. In this regard it is interesting to think of the functional relationship between these two behavioural rhythms. The exploration of a territory and the exploitation of its resources are complementary: the exploitation of known resources provides predictable benefits, whereas exploration reduces uncertainty about the availability of unknown resources in other areas of the territory (Mehlhorn et al., 2015; Petzke and Schomaker, 2022; Reader, 2015). From a locomotor perspective, exploration is characterized by the displacement of the individual across the territory, whereas exploitation involves a longer stay at the resource site with locomotor activity that does not imply displacement (Mehlhorn et al., 2015). The nocturnal increase in EODr enhances the amount of sensory information available and we can propose two possible functions for this nocturnal increase: first, one that is independent of displacement and possibly linked to the exploitation of nearby resources, and second, one associated with patrolling and exploring the territory. In this sense, it is important to note that our analysis will underestimate movements whose trajectories are shorter than 20 cm (the approximate length of an adult *G. omarorum*). The persistence of exploratory activity in the absence of social stimuli supports the view that exploration in *G. omarorum* is an intrinsic, self-sustained behavior likely associated with territory maintenance and feeding. The preference for peripheral regions of the arena suggests that fish engage in patrol-like movements that mimic boundary exploration, similar to the nocturnal territorial patrolling observed in field studies (Migliaro et al., 2005).

Our findings position G. omarorum as a valuable system for examining how animals organize sensory sampling and movement across temporal and environmental contexts. By demonstrating that exploratory behavior emerges from the coordinated modulation of locomotor and electric behavior rhythms, our study links mechanistic aspects of sensory physiology, chronobiology and ecologically relevant patterns of territory use. Although much of the work on motivated exploration has relied on model organisms in highly controlled settings (Gordon et al., 2014; Haley and Chalasani, 2024), our results highlight the importance of incorporating semi-natural protocols that preserve environmental structure while allowing selective experimental control (Flôres et al., 2021; Sanguinetti-Scheck and Gálvez, 2024; Shemesh et al., 2024). This integrative approach reveals consistent individual differences in the timing of electric behavior and locomotor activity, opening new opportunities to investigate wild chronotypes, individual strategies of territory use, and the neural mechanisms coupling electrosensory sampling to movement.

## Acknowledgements

Wéd like to thank Jan Benda for kindly sharing the teensys and amplifiers we used in the recording grids. We thank Cecilia Jalabert, Guillermo Valiño, Rossana Perrone, María Victoria Falco and Diego Gallo for their help in fish collection and semi-natural experiments. We are also grateful to Ana Silva who read and commented on an earlier version of our manuscript.

## Financial support

CAP_Udelar. Universidad de la República, Uruguay.

ANII_FCE_1_2021_1_167077. Agencia Nacional de Investigación e Innovación. Uruguay. Udelar_ Grupo CSIC 883. Modulación ambiental y social del reloj biológico. Universidad de la República, Uruguay.

## Data and resource availability

