## Supplementary material for "One More Lap: Environmental Modulation of Small-Scale Exploration in Weakly Electric Fish": Supp Material

Data is available upon request and will be available on a repository by the moment of publication.

### Diversity and inclusion statement

The authors recognize that diversity and inclusion strengthen scientific research and are committed to equitable and inclusive practices in their collaborative, experimental, and dissemination activities.

### Supplementary material

| Group 1 | Group 2 | Wilcoxon pair-wise test | | |
|---|---|---|---|---|
| | | W+ | Uncorrected p-value | Adjusted p-value (Bonferroni-Holm) |
| ***XD*** | ***XL*** | ***0*** | ***0.0312**** | ***0.4688*** |
| XD | SD(E) | 1 | 0.0625 | 0.6875 |
| XD | SD(M) | 10 | 1 | 1 |
| XD | SL(E) | 10 | 1 | 1 |
| XD | SL(M) | 7 | 0.5625 | 1 |
| ***XL*** | ***SD(E)*** | ***0*** | ***0.0312**** | ***0.4375*** |
| ***XL*** | ***SD(M)*** | ***0*** | ***0.0312**** | ***0.4062*** |
| XL | SL(E) | 0 | 0.1088 | 1 |
| **XL** | **SL(M)** | **0** | **0.0431*** | **0.5174** |
| SD(E) | SD(M) | 6 | 0.4375 | 1 |
| SD(E) | SL(E) | 6 | 0.4375 | 1 |
| SD(E) | SL(M) | 5 | 0.3125 | 1 |
| SD(M) | SL(E) | 10 | 1 | 1 |
| SD(M) | SL(M) | 6 | 0.4375 | 1 |
| SL(E) | SL(M) | 5 | 0.5002 | 1 |

**Table S1: Statistical results of pair-wise comparisons of locomotor activity during different contexts in laboratory experiments.** For each pair of condition groups, the table reports the Wilcoxon test statistic (W+), the uncorrected p-value, and the Holm–Bonferroni adjusted p-value for multiple comparisons. Asterisks (*) denote uncorrected p-values below 0.05. XD=stimulus Off/Darkness, XL =stimulus On/Light, SD(E) = Stimulus on/Darkness (Evening trial), SD(M) = Stimulus On/Darkness (Morning trial), SL(E) = Stimulus On/Light (Evening trial), SL(M) = Stimulus On/Light (Morning trial).

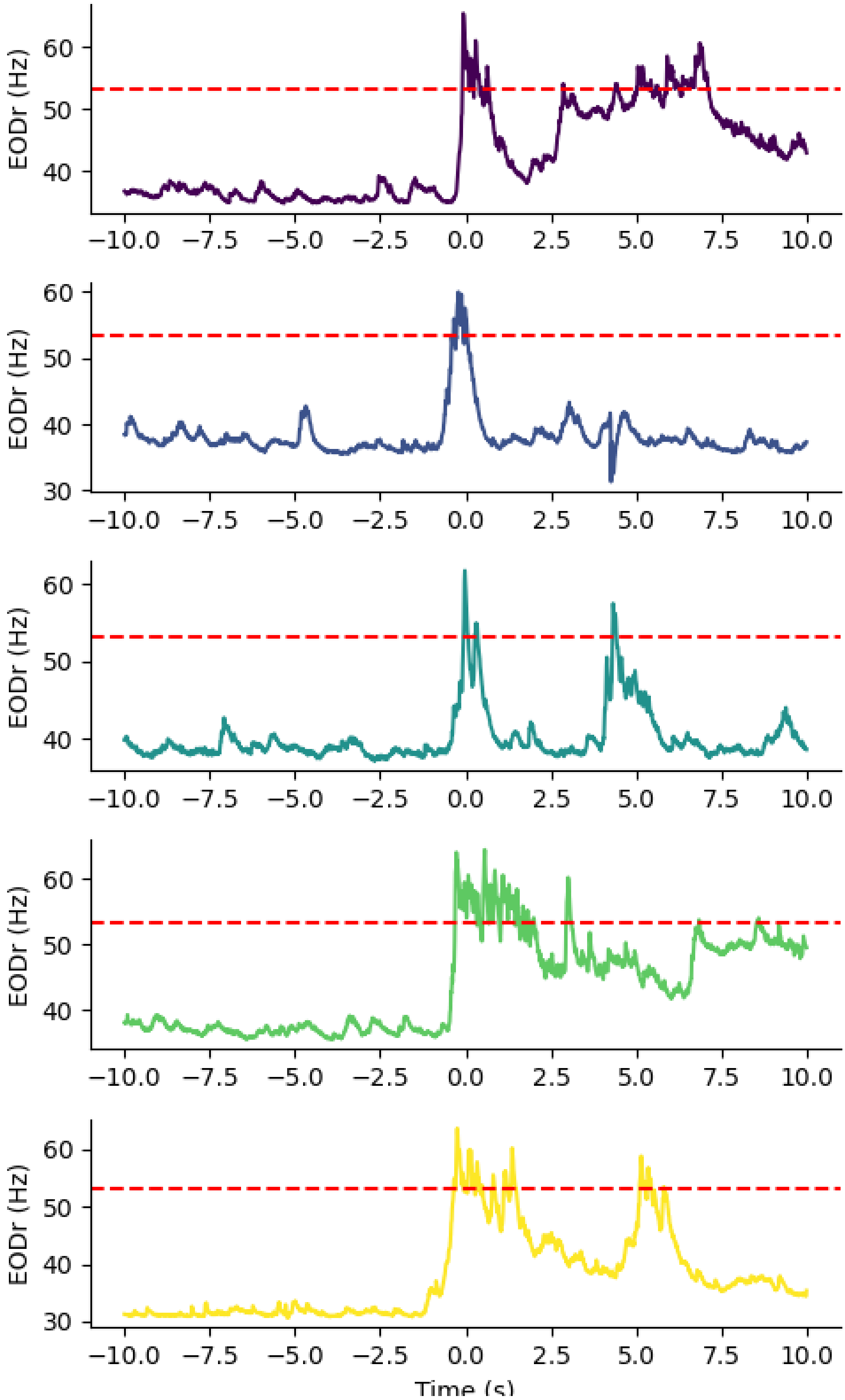


**Figure S1. Examples of high-frequency events (HFEs) in a single fish.** Each panel shows the EOD rate (EODr, in Hz) over a 20-second window, centered on the occurrence of a high-frequency event (time 0 s). The red dashed line indicates the threshold for HFE detection (1.5×z-score). To qualify as a high-frequency event, the EODr had to exceed this threshold for at least three consecutive EODs.

| Group 1 | Group 2 | Wilcoxon pair-wise test | | |
|---|---|---|---|---|
| | | **W+** | **Uncorrected p-value** | **Adjusted p-value (Bonferroni-Holm)** |
| XL | XD | 6 | 0.4375 | 0.875 |
| XL | SL(E) | 2 | 0.0938 | 0.6562 |
| XL | SD(E) | 1 | 0.0625 | 0.625 |
| ***XL*** | ***SL(M)*** | ***0*** | ***0.0312**** | ***0.4688*** |
| ***XL*** | ***SD(M)*** | ***0*** | ***0.0312**** | ***0.4375*** |
| XD | SL(E) | 1 | 0.0625 | 0.5625 |
| XD | SD(E) | 0 | 0.0312 | 0.4062 |
| XD | SL(M) | 2 | 0.0938 | 0.5625 |
| ***XD*** | ***SD(M)*** | ***0*** | ***0.0312**** | ***0.375*** |
| *SL(E)* | *SD(E)* | *3* | *0.1562* | *0.625* |
| SL(E) | SL(M) | 5 | 0.3125 | 0.9375 |
| ***SL(E)*** | ***SD(M)*** | ***0*** | ***0.0312**** | ***0.3438*** |
| SD(E) | SL(M) | 9 | 0.8438 | 0.8438 |
| SD(E) | SD(M) | 2 | 0.0938 | 0.4688 |
| SL(M) | SD(M) | 1 | 0.0625 | 0.5 |

**Table S2: Statistical results of pair-wise comparison of the electromotor response index during different contexts in laboratory experiments.** For each pair of condition groups, the table reports the Wilcoxon test statistic (W+), the uncorrected p-value, and the Holm–Bonferroni adjusted p-value for multiple comparisons. Asterisks (*) denote uncorrected p-values below 0.05. XD=stimulus Off/Darkness, XL =stimulus On/Light, SD(E) = Stimulus on/Darkness (Evening trial), SD(M) = Stimulus On/Darkness (Morning trial), SL(E) = Stimulus On/Light (Evening trial), SL(M) = Stimulus On/Light (Morning trial).

| Fish | p | q | Adjusted R2 | Amplitude (%) | Acrophase (rad) | Acrophase (hs) | MESOR (%) |
|---|---|---|---|---|---|---|---|
| **1** | **0.052** | **0.052** | **0.01** | **1.28** | **-1.70** | **6.49** | **8.56** |
| 2 | <0.0001 | <0.0001 | 0.35 | 9.28 | -0.16 | 0.62 | 18.68 |
| 3 | <0.0001 | <0.0001 | 0.31 | 7.63 | 0.07 | 23.74 | 9.73 |
| 4 | <0.0001 | <0.0001 | 0.45 | 9.30 | -0.36 | 1.39 | 12.50 |
| 5 | <0.0001 | <0.0001 | 0.38 | 17.70 | -0.16 | 0.62 | 21.56 |
| 6 | <0.0001 | <0.0001 | 0.30 | 5.58 | 0.12 | 23.54 | 4.90 |
| 7 | <0.0001 | <0.0001 | 0.38 | 7.77 | -0.06 | 0.24 | 8.72 |
| 8 | <0.0001 | <0.0001 | 0.22 | 5.88 | 0.36 | 22.63 | 13.84 |
| 9 | <0.0001 | <0.0001 | 0.27 | 7.09 | 0.06 | 23.78 | 8.64 |

**Table S3: Cosinor analysis parameters for daily rhythmicity in locomotor activity.** For each fish, the table shows the significance of the cosine fit (p) and the FDR-corrected significance (q), the proportion of variance explained (Adjusted $R^2$), and the rhythmic parameters: Amplitude (deviation from the MESOR), Acrophase (timing of model peak, in radians and converted to hours), and MESOR (midline estimating statistic of rhythm, expressed as percentage). Higher amplitudes indicate stronger rhythmicity, while the acrophase reflects the time of day at which the variable reaches its maximum.

| Fish | p | q | Adjusted R2 | Amplitude (Hz) | Acrophase (rad) | Acrophase (hs) | MESOR (Hz) |
|---|---|---|---|---|---|---|---|
| 1 | <0.0001 | <0.0001 | 0.50 | 5.42 | 0.50 | 22.10 | 35.14 |
| 2 | <0.0001 | <0.0001 | 0.45 | 4.11 | 0.32 | 22.77 | 33.85 |
| 3 | <0.0001 | <0.0001 | 0.66 | 5.95 | 0.55 | 21.91 | 31.55 |
| 4 | <0.0001 | <0.0001 | 0.55 | 4.36 | -0.05 | 0.19 | 32.42 |
| 5 | <0.0001 | <0.0001 | 0.39 | 4.41 | 0.02 | 23.93 | 30.52 |
| 6 | <0.0001 | <0.0001 | 0.24 | 3.09 | 1.60 | 17.87 | 27.06 |
| 7 | <0.0001 | <0.0001 | 0.34 | 3.33 | 0.67 | 21.43 | 33.60 |
| 8 | <0.0001 | <0.0001 | 0.52 | 4.22 | 0.84 | 20.80 | 31.62 |
| 9 | <0.0001 | <0.0001 | 0.43 | 4.49 | 0.70 | 21.33 | 31.44 |

**Table S4: Cosinor analysis parameters for daily rhythmicity in EOD rate.** For each fish, the table shows the significance of the cosine fit (p) and the FDR-corrected significance (q), the proportion of variance explained (Adjusted $R^2$), and the rhythmic parameters: Amplitude (deviation from the MESOR), Acrophase (timing of model peak, in radians and converted to hours), and MESOR (midline estimating statistic of rhythm, expressed in Hz).
